# Pharmacological rescue of impaired mitophagy in Parkinson’s disease-related LRRK2 G2019S knock-in mice

**DOI:** 10.1101/2020.12.07.414359

**Authors:** Francois Singh, Alan R. Prescott, Graeme Ball, Alastair D. Reith, Ian G. Ganley

## Abstract

Parkinson’s disease (PD) is a major and progressive neurodegenerative disorder, yet the biological mechanisms involved in its aetiology are poorly understood. Evidence links this disorder with mitochondrial dysfunction and/or impaired lysosomal degradation – key features of the autophagy of mitochondria, known as mitophagy. Here we investigated the role of LRRK2, a protein kinase frequently mutated in PD, on this process *in vivo*. Using mitophagy and autophagy reporter mice, bearing either knockout of LRRK2 or expressing the pathogenic kinase-activating G2019S LRRK2 mutation, we found that basal mitophagy was specifically altered in clinically relevant cells and tissues. Our data show that basal mitophagy inversely correlates with LRRK2 kinase activity *in vivo*. In support of this, use of distinct LRRK2 kinase inhibitors in cells increased basal mitophagy, and a CNS penetrant LRRK2 kinase inhibitor, GSK3357679A, rescued the mitophagy defects observed in LRRK2 G2019S mice. This study provides the first *in vivo* evidence that pathogenic LRRK2 directly impairs basal mitophagy, a process with strong links to idiopathic Parkinson’s disease, and demonstrates that pharmacological inhibition of LRRK2 is a rational mitophagy-rescue approach and potential PD therapy.

## Introduction

Parkinson’s disease (PD) is the second most common neurodegenerative disorder, affecting 1-2% of the population over 60 years old, and 4% above 85 years of age ^1^. The main symptoms of PD are muscle rigidity, bradykinesia, resting tremor, and postural instability and may be accompanied by sleep disorders, anosmia, depression, and dementia ^2^. It is characterised by a progressive and selective degeneration of dopaminergic (DA) neurons of the substantia nigra pars compacta (SNpc). Currently, there are no treatments available that modify the course of neurodegenerative decline. Although this disease is mostly sporadic, about 15% of cases appear to be inherited and in support of this, 20 genes implicated in PD have been identified from familial genetic studies while ~90 loci have been identified from PD-GWAS ^3^. The exact causes of PD are currently unknown but some evidence strongly links impaired mitochondrial and lysosomal function to disease pathology ^2^.

Mutations in *PARK8*, encoding for LRRK2 (Leucine Rich Repeat Kinase 2), are the most frequently reported cause of PD ^4^. The most common mutation associated with PD is the substitution of glycine at position 2019 of LRRK2 to serine (G2019S), representing 4% of familial and 1% of sporadic cases ^5^. LRRK2 is a large multidomain protein with two catalytic domains: a Ras of complex (ROC) GTPase domain that is able to bind GTP and hydrolyse it, and a kinase domain that utilises a subset of Rab GTPases as substrates ^6^. Importantly, all the segregating mutations associated with PD are located in the catalytic core. A mutation in the ROC/COR domain, such as the R1441C/G/H or the Y1699C mutation, leads to decreased GTPase activity and elevated kinase activity ^7^. Mutations in the kinase domain, such as the G2019S or the I2020T, also lead to an elevated kinase activity. Hence, enhanced kinase activity appears to be a common factor in pathogenic LRRK2 mutations. Although the function of LRRK2 within cells is currently unknown, mounting evidence implicates a role in membrane trafficking ^6,8,9^.

Macroautophagy is a membrane trafficking pathway that delivers intracellular components to the lysosome for degradation ^10^. These components can include whole organelles such as mitochondria. The autophagic turnover of mitochondria is termed mitophagy, which acts as a mitochondrial quality control mechanism that allows the selective degradation of damaged or unnecessary mitochondria ^11,12^. Mitophagy itself has strong links to PD following the landmark discoveries that PINK1 and Parkin, two other genes mutated in familial PD, sequentially operate to initiate mitophagy in response to mitochondrial depolarisation in cell lines ^13–16^. However, when this pathway becomes relevant *in vivo*, and under what physiological conditions, is unclear especially given that PINK1 and Parkin are not required for regulation of mitophagy under normal, or basal, conditions ^17–19^. Indeed, our understanding of the detailed mechanisms regulating basal mitophagy remains elusive.

In this study, we sought to define the physiological link between mitochondrial turnover and LRRK2 in relation to PD. We utilised our previously published mouse reporter models to study mitophagy (*mito*-QC) and autophagy (*auto*-QC; Fig. 1A, C and ^17,20,21^) in either LRRK2 knockout mice, or knock-in mice harbouring the pathogenic LRRK2 G2019S mutation. Whilst we found minimal impact of LRRK2 on general autophagy (macroautophagy), we observed that the LRRK2 G2019S activation-mutation was associated with reduced mitophagy in specific tissues, including dopaminergic neurons and microglia within the brain. In contrast, knockout of LRRK2 resulted in increased mitophagy. Taken together, these data imply that LRRK2 kinase activity inversely correlates with basal mitophagy levels. In support of this, we found that that treatment of cells or animals with the potent and selective CNS penetrant LRRK2 kinase inhibitor, GSK3357679A (Ding et al., in preparation), rescued these LRRK2 G2019S-associated mitophagy defects and enhanced mitophagy in dopamine neurons and microglia in the brains of genotypically normal mice. Our results identify a physiological role for LRRK2 in the regulation of basal mitophagy *in vivo* and underline the potential value of pharmacological inhibition of LRRK2 as a potential therapeutic strategy to ameliorate aspects of Parkinson’s disease driven by mitochondrial dysfunction.

**Figure 1.**
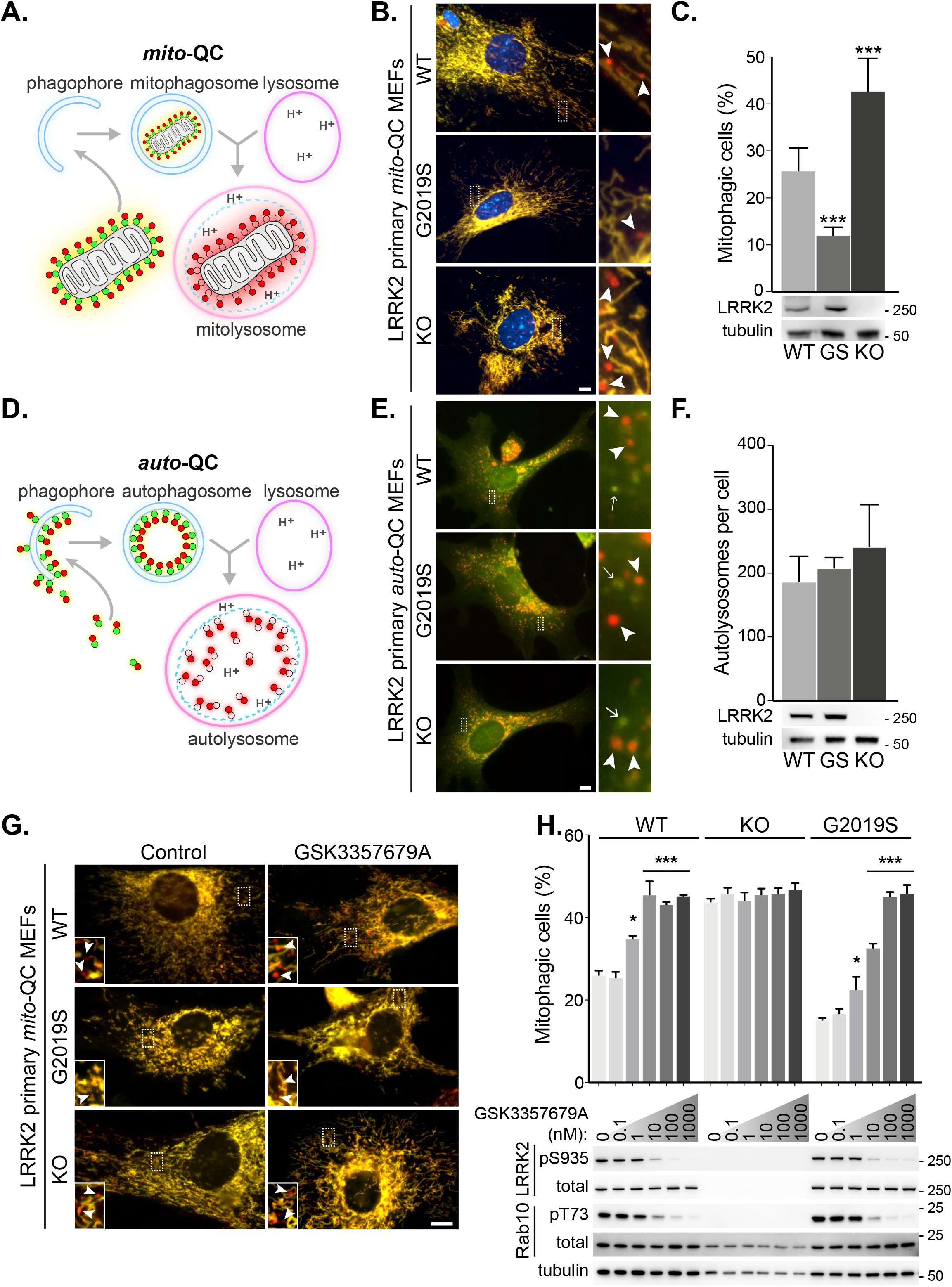
LRRK2 kinase activity impairs basal mitophagy *in vitro*. (**A**) Schematics of the *mito*-QC reporter in mouse model. (**B**) Representative images of *mito*-QC primary MEF cultures established from LRRK2 WT, LRRK2 G2019S, and LRRK2 KO embryos. Boxed area is magnified on the right and arrowheads indicate examples of mitophagy (mCherry-only mitolysosomes). (**C**) Quantitation of data shown in B from 6-9 independent experiments. Below is representative immunoblot showing LRRK2 protein expression. (**D**) Schematics of the *auto*-QC reporter in mouse model. (**E**) Representative images of *auto*-QC primary MEF cultures established from LRRK2 WT, LRRK2 G2019S, and LRRK2 KO embryos. Boxed area is magnified on the right and arrowheads indicate examples of autolysosomes and arrows highlight autophagosomes. (**F**) Quantitation of data shown in E from 4-6 independent experiments. (**G**) Representative images of *mito*-QC primary MEFs treated with control (DMSO) or 100 nM GSK3357679A. Boxed area is magnified on bottom left and arrowheads indicate examples of mitolysosomes. (**H**) Quantitation of mitophagy shown in G from 3-7 independent experiments. Corresponding immunoblot of indicated proteins is shown below. Scale bars, 10 μm. Overall data is represented as mean +/−SEM. Statistical significance is displayed as *p<0.05, and ***p<0.001.

## Material and methods

### Animals

Experiments were performed on mice genetically altered for Leucine-rich repeat kinase 2 (LRRK2), using either wild-type, LRRK2 G2019S ^6^ mutation knock-in mice, or mice in which LRRK2 has been ablated ^22,23^ (KO).The mitophagy (*mito*-QC) and the autophagy (*auto*-QC) reporter mouse models used in this study were generated as previously described ^17,20^.

### Primary mouse embryonic fibroblasts culture

Primary mouse embryonic fibroblasts (MEFs) were derived, from time-mated pregnant females at E12.5. Primary MEFs were maintained in DMEM (Gibco, 11960-044) supplemented with 10% FBS, 2 mM L-Glutamine (Gibco, 2503-081), 1% Na-Pyruvate (Gibco, 11360-070), 1% Non-essential amino acids (Gibco, 11140-035), 1% Antibiotics (Penicillin/Streptomycin 100 U/ml penicillin and 100 μg/ml streptomycin; Gibco), and 150μM β-Mercaptoethanol (Gibco, 21985-023) at 37°C under a humidified 5% CO_2_ atmosphere.

### Primary mouse embryonic fibroblasts treatments

To assess mitophagy and autophagy upon stimulation, cells were treated for 24 hours with either 1 mM 3-Hydroxy-1,2-dimethyl-4(1H)-pyridone (Deferiprone/DFP, Sigma-Aldrich, 379409), or incubated in Earl’s balanced salt solution (EBSS, Gibco, 24010-043). mito-QC MEFs were also treated for 24 hours with LRRK2 kinase activity inhibitors GSK2578215A ^24^ (250, 500, 1000 nM)), MLi-2 ^25^ (5, 10, and 20 nM), or GSK3357679A (compound **39**, Ding et al., in prep.) (0.1, 1, 10, 100, 1000 nM). All treatments (apart from EBSS) were in DMEM (Gibco, 11960-044) supplemented with 10% FBS, 2 mM L-Glutamine (Gibco, 2503-081), 1% Non-essential amino acids (Gibco, 11140-035), 1% Antibiotics (Penicillin/Streptomycin 100 U/ml penicillin and 100 μg/ml streptomycin; Gibco), and 150μM β-Mercaptoethanol (Gibco, 21985-023) at 37°C under a humidified 5% CO_2_ atmosphere. MLi-2 and GSK2578215A were synthesized by Natalia Shapiro (University of Dundee) as described previously ^24,25^.

### Light Microscopy

MEFs were plated on glass coverslips and treated as described in the previous paragraph. At the end of the treatment, cells were washed twice in DPBS (Gibco, 14190-094), and fixed in 3.7% Paraformaldehyde (Sigma, P6148), 200 mM HEPES, pH=7.00 for 20 minutes. Cells were washed twice with, and then incubated for 10 minutes with DMEM, 10mM HEPES. After a wash with DPBS, nuclei were stained with Hoechst 33342 (1 μg/mL, Thermo Scientific, 62249) for 5 minutes. Cells were washed in DPBS and mounted on a slide (VWR, Superfrost, 631-0909) with Prolong Diamond (Thermo Fisher Scientific, P36961). Images were acquired using a Nikon Eclipse Ti-S fluorescence microscope with a 63x objective.

### Quantitation of Mitophagy and Autophagy in vitro

Quantification of red-only dots was semi-automatized using the *mito*-QC counter plugin on FIJI as previously described ^26,27^. Autophagosomes and autolysosomes were quantified using the Autophagy counter plugin on FIJI developed in house, following the same principle as the mito-QC counter ^28^. The macro “auto-QC_counter.ijm” (“version 1.0 release”, DOI: 10.5281/zenodo.4158361) is available from the following github repository: https://github.com/graemeball/auto-QC_counter.

### Animal Studies

Experiments were performed on 81 adult mice (9-23 weeks old) of both genders (n=8-12 per group for the mito-QC reporter, and n=9-15 per group for the auto-QC reporter), all homozygous for the corresponding reporter (mitophagy or autophagy).

The effect of the CNS penetrant LRRK2 kinase inhibitor GSK3357679A *in vivo* was assessed using 50 adult mice (9-17 weeks old at the end of the study), all homozygous for the mito-QC reporter. Mice of both genders were randomly assigned to the vehicle or to the GSK3357679A treated group (WT-Vehicle: n=10, WT-GSK3357679A: n=10, G2019S-Vehicle: n=10, G2019S-GSK3357679A: n=10, KO-Vehicle: n=5, and KO-GSK3357679A: n=5). Vehicle treated animals were dosed (10 mL/kg) with aqueous methylcellulose (1% w/v, Sigma, M0512) prepared in sterile water (Baxter, UKF7114), or with GSK3357679A (15 mg/kg/dose) prepared in aqueous methylcellulose. Treatment was administered by oral gavage every 12 hours for a total of four times per mouse. Mice were culled 2 hours (+/−9 minutes) after the last dosing.

Animals were housed in sex-matched littermate groups of between two and five animals per cage in neutral temperature environment (21° ± 1 °C), with a relative humidity of 55-65%, on a 12:12 hour photoperiod, and were provided food and water *ad libitum*. All animal studies were ethically reviewed and carried out in accordance with Animals (Scientific Procedures) Act 1986 as well as the GSK Policy on the Care, Welfare and Treatment of Animals, and were performed in agreement with the guidelines from Directive 2010/63/EU of the European Parliament on the protection of animals used for scientific purposes. All animal studies and breeding were approved by the University of Dundee ethical review committee, and further subjected to approved study plans by the Named Veterinary Surgeon and Compliance Officer (Dr. Ngaire Dennison) and performed under a UK Home Office project license in agreement with the Animal Scientific Procedures Act (ASPA, 1986).

### Sample collection

Mice were terminally anesthetised with an intraperitoneal injection of pentobarbital sodium (Euthatal, Merial) then trans-cardially perfused with DPBS (Gibco, 14190-094) to remove blood. Tissues were collected and either snap frozen in liquid nitrogen and stored at −80°C for later biochemical analyses or processed by overnight immersion in freshly prepared fixative: 3.7% Paraformaldehyde (Sigma, P6148), 200 mM HEPES, pH=7.00. The next day, fixed tissues were washed three times in DPBS, and immersed in a sucrose 30% (w/v) solution containing 0.04% sodium azide until they sank at the bottom of the tube. Samples were stored at 4°C in that sucrose solution until further processing.

### Immunolabeling of brain free-floating sections

The brain was frozen-sectioned axially using a sledge microtome (Leica, SM2010R), and 50 microns thick sections were stored in PBS at 4°C until further treatment. Free-floating sections were permeabilised using DPBS (Gibco, 14190-094) containing 0.3% Triton X-100 (Sigma Aldrich, T8787) 3 times for 5 minutes. Sections were then blocked for one hour in blocking solution (DPBS containing 10% goat serum (Sigma Aldrich, G9023), and 0.3% Triton X-100). Primary antibody incubation was performed overnight in blocking solution containing one of the following antibodies: Anti-Tyrosine Hydroxylase (1/1000, Millipore, AB152), Anti-Iba-1 (1/1000, Wako, 019-19741), Anti Calbindin-D28k (1/1000, Swant, CB38), Anti-Glial Fibrillary Acidic Protein (1/1000, Millipore, MAB360). The next day, sections were washed 2 times for 8 minutes in DPBS containing 0.3% Triton X100 and then incubated for 1 hour in blocking solution containing the secondary antibody (1/200, Invitrogen P10994 Goat anti-Rabbit IgG (H+L) Cross-Adsorbed Secondary Antibody, Pacific Blue, or Invitrogen P3182 Goat anti-Mouse IgG (H+L) Cross-Adsorbed Secondary Antibody, Pacific Blue). Sections were then washed 2 times for 8 minutes in DPBS containing 0.3% Triton X100 and mounted on slides (Leica Surgipath^®^ X-tra™ Adhesive, 3800202) using Vectashield Antifade Mounting Medium (Vector Laboratories, H-1000) and sealed with nail polish.

### Tissue section and immunostaining

Tissues were embedded in an O.C.T. compound matrix (Scigen, 4586), frozen and sectioned with a cryostat (Leica CM1860UV). 12 microns sections were placed on slides (Leica Surgipath^®^ X-tra™ Adhesive, 3800202), and then air dried and kept at −80°C until further processing. Sections were thawed at room temperature and washed 3 times 5 minutes in DPBS (Gibco, 14190-094). Sections were then counterstained for 5 minutes with Hoechst 33342 (1μg/mL, Thermo Scientific, 62249). Slides were mounted using Vectashield Antifade Mounting Medium (Vector Laboratories, H-1000) and high precision cover glasses (No. 1.5H, Marienfeld, 0107222) and sealed with transparent nail polish.

### Confocal microscopy

Confocal micrographs were obtained by uniform random sampling using either a Zeiss LSM880 with Airyscan, or a Zeiss 710 laser scanning confocal microscope (Plan-Apochromat 63x/1.4 Oil DIC M27) using the optimal parameters for acquisition (Nyquist). 10-15 images were acquired per sample, depending on the tissue, by an experimenter blind to all conditions. High resolution, representative images were obtained using the Super Resolution mode of the Zeiss LSM880 with Airyscan.

### Quantitation of Mitophagy and Autophagy in vivo

Quantification of mitophagy and autophagy was carried out on at least 10 pictures per sample. Images were processed with Volocity Software (version 6.3, Perkin-Elmer). Images were first filtered using a fine filter to suppress noise. Tissue was detected by thresholding the Green channel. For the immunolabelings in the brain (TH, Iba1, Calbindin D-28k, and GFAP), each cell population of interest was detected by thresholding the Pacific Blue labelled channel. A ratio image of the Red/Green channels was then created for each image. For the mito-QC reporter, mitolysosomes were then detected by thresholding the ratio channel as objects with a high Red/Green ratio value within the tissue/cell population of interest. The same ratio channel threshold was used per organ/set of experiments. To avoid the detection of unspecific high ratio pixels in the areas of low reporter expression, a second red threshold was applied to these high ratio pixels. This double thresholding method provides a reliable detection of mitolysosomes as structures with a high Red/Green ratio value and a high Red intensity value.

For the general autophagy reporter, high intensity red pixels were detected by thresholding the red channel within the tissue/cell population of interest. The same red channel threshold was used per organ/set of experiments. Autophagosomes and autolysosomes were then differentiated by thresholding the high intensity red pixels depending on their Red/Green ratio channel value. Pixels with a low Red/Green ratio were considered as autophagosomes, whereas pixels with a high Red/Green ratio were considered as autolysosomes. The same ratio channel threshold was used per organ/set of experiments.

### Filipin staining

Frozen fixed tissue sections were washed in PBS to remove any excess O.C.T compound (Scigen, 4586) excess. Sections were then incubated for 2 hours at room temperature with filipin (200 μg/mL; Sigma-Aldrich, F9765) and then washed twice in PBS. Tissue sections were mounted using Vectashield Antifade Mounting Medium (Vector Laboratories, H-1000) and sealed with nail polish. High resolution, representative images were obtained using the Super Resolution mode of the Zeiss LSM880 with Airyscan (Plan-Apochromat 63x/1.4 Oil DIC M27).

### Western blotting

Frozen tissue was homogenized with a Cellcrusher™ (Cellcrusher, Cork, Ireland) tissue pulveriser. Approximately 20-30 mg of pulverised tissue were then lysed on ice for 30 min with (10 μL/mg tissue) of RIPA buffer [50 mM Tris–HCl pH 8, 150 mM NaCl, 1 mM EDTA, 1% NP-40, 1% Na-deoxycholate, 0.1% SDS, and cOmplete™ protease inhibitor cocktail (Roche, Basel, Switzerland)], phosphatase inhibitor cocktail (1.15 mM sodium molybdate, 4 mM sodium tartrate dihydrate, 10 mM β-glycerophosphoric acid disodium salt pentahydrate, 1 mM sodium fluoride, and 1mM activated sodium orthovanadate), and 10 mM DTT. After lysis, the mixture was vortexed and centrifuged for 10 min at 4 °C at 20,817 G. The supernatant was collected, and the protein concentration determined using the Pierce BCA protein assay kit (ThermoFisher Scientific, Waltham, MA, USA). For each sample, 20-25 μg of protein was separated on a NuPAGE 4–12% Bis-Tris gel (Life technologies, Carlsbad, CA, USA). Proteins were electroblotted to 0.45μm PVDF membranes (Imobilon-P, Merck Millipore, IPVH00010; or Amersham Hybond, GE Healthcare Life Science, 10600023), and immunodetected using primary antibodies directed against phospho-Ser935 LRRK2 rabbit monoclonal (1/1000, MRC PPU Reagents and Services, UDD2), LRRK2 rabbit monoclonal (1/1000, MRC PPU Reagents and Services, UDD3), phospho-Rab10 (Thr73) rabbit monoclonal (1/1000, Abcam, ab230261), Rab10 mouse monoclonal (1/1000, nanoTools 0680-100/Rab10-605B11), α-Tubulin (11H10) Rabbit monoclonal antibody (1/10000, CST, 2125S), and β-Actin mouse monoclonal antibody (1/1000, Proteintech, 60008-1-Ig). All antibodies to LRRK2 were generated by MRC PPU Reagents and Services, University of Dundee (http://mrcppureagents.dundee.ac.uk).

### Statistics

Data are represented as means ± SEM. Number of subjects are indicated in the respective figure legends. Statistical analyses were performed using a one-way analysis of variance (ANOVA) or two-way ANOVA followed by a Tukey HSD using RStudio version 1.1.1335 ^29^. Statistical significance is displayed as * p< 0.05: ** p < 0.01, *** p<0.001, and **** p<0.0001.

## Results

### *The pathogenic G2019S LRRK2 mutation impairs basal mitophagy but not autophagy* in vitro

To investigate the physiological role of LRRK2 in regulating autophagy we utilised two previously validated and highly similar mouse reporter models ^17,20,21^. These transgenic reporter models rely on constitutive expression of a tandem mCherry-GFP tag from the *Rosa26* locus. In the *mito*-QC model, which monitors mitophagy, the tandem tag is localised to mitochondria (by an outer mitochondrial targeting sequence derived from residues 101-152 of the protein FIS1). In the *auto*-QC model, which monitors general (macro)autophagy, the tandem tag is localised to autophagosomes (by conjugation to the N-terminus of MAP1LC3b). For both models, when a mitochondrion or autophagosome is delivered to lysosomes, the low lysosomal luminal pH is sufficient to quench the GFP signal, but not that from mCherry. Hence, the degree of mitophagy or general autophagy can be determined by the appearance of mCherry-only puncta, which represent mito/autolysosomes (Fig. 1A and D). Given that mitophagy is a form of autophagy, the use of both models allows us to monitor the specificity of autophagy *in vivo*. A large disruption of autophagy in general will also influence mitophagy, whereas a block in mitophagy, which likely represents a small fraction of the total autophagy occurring at any one time, will tend to have little influence on the total autophagic levels.

To investigate the effect of LRRK2 kinase activity on mitophagy, we first isolated and cultured primary mouse embryonic fibroblasts (MEFs) derived from wild type mice (WT), mice homozygous for the Parkinson’s disease-associated LRRK2 G2019S mutant, or mice homozygous for a LRRK2 knockout (KO) variant; all of which were on a homozygous *mito*-QC reporter background (Fig. 1B and C). A small degree of basal mitophagy was evident in all cell lines. However, we observed that LRRK2 G2019S KI mutant cells displayed significantly lower basal mitophagy levels, whereas the absence of LRRK2 (KO) led to an increase of this process. Interestingly, our data suggests that LRRK2 predominantly influences basal mitophagy as deferiprone (DFP), a strong mitophagy inducer ^30^, increased mitophagy to a similar level across all genotypes (Fig. S1A and B).

We next investigated general autophagy using the LRRK2 mouse lines mentioned above on the homozygous *auto*-QC background. In contrast to mitophagy, in isolated primary MEFs we noticed no significant difference in the number of mCherry-only autolysosomes across all the *Lrrk2* genotypes under basal conditions (Fig. 1E and F). We also analysed amino acid starvation-induced autophagy, by incubation in Earls Balanced Salt Solution (EBSS). A robust autophagy response was observed in all cells and as with basal autophagy, the *Lrrk2* genotype failed to significantly alter this large increase in autolysosomes (Fig. S1C).

### LRRK2 kinase inhibitors correct the G2019S mitophagy defects in vitro

Using genetics, our observations show that LRRK2 kinase activity inversely correlates with mitophagy *in vitro*. If this is the case, then pharmacological inhibition of LRRK2 kinase activity should also increase mitophagy. Therefore, we aimed to investigate if the mitophagy deficit observed in the G2019S cells could be rescued with LRRK2-selective kinase inhibitors. To that end, we first studied the effect of two structurally distinct tool LRRK2 kinase inhibitors, GSK2578215A ^24^ and MLi-2 ^25^, in primary *mito*-QC MEFs. In the WT group, treatment of cells for 24 h with either compound increased mitophagy to values comparable to what we previously observed in the LRRK2 KO group (Fig. S1D). Although MLi-2 is a more potent LRRK2 kinase inhibitor (IC_50_ 0.76 nM ^25^) compared to GSK2578215A (IC_50_ 10nM^24^), at higher concentrations it did not stimulate mitophagy. In the G2019S group, significantly higher concentrations of GSK2578215A were necessary to stimulate mitophagy and this may reflect the increased kinase activity of this mutant ^31^. However, with MLi-2 we were not able to fully rescue the mitophagy defect. As with the WT cells, treatment with MLi-2 at the highest dose (20 nM) did not stimulate mitophagy. The failure of higher doses of MLi-2 to stimulate mitophagy may be due to an off-target effect, as at 20 nM it also inhibited mitophagy in the LRRK2 KO cells (Fig. S1D). Therefore, we recommend caution when using MLi-2 at high concentrations. Neither compound increased mitophagy in KO cells, demonstrating that their mitophagy-enhancing properties are dependent on LRRK2. Additionally, both compounds inhibited LRRK2 activity, as indicated by loss of LRRK2 S935 phosphorylation (an indirect measure of LRRK2 activity ^32^, Fig. S1E).

With respect to mitophagy, the relatively low potency of GSK2578215A and the off-target effects of MLi2, indicated the need for improved LRRK2 tool inhibitors. For this reason, we turned to GSK3357679A, a novel pyrrolopyrimidine LRRK2 kinase inhibitor that exhibits excellent cellular potency, selectivity, oral bioavailability and PK/PD correlation in animal studies (Ding et al., in prep.). We tested GSK3357679A in primary *mito*-QC MEFs and observed a dose-dependent effect on mitophagy with a maximal stimulation achieved at a concentration of 10 nM in WT cells (Fig. 1G and H). In the G2019S cells we observed a reduced response, with a maximal effect on mitophagy reached at 100 nM. Importantly, contrary to what we observed with MLi-2, GSK3357679A effects on restoration of mitophagy were maintained at the highest concentration used and exhibited no deleterious effects on mitophagy at any concentration in LRRK2 KO cells. Western blotting analysis revealed that GSK3357679A potently inhibited LRRK2 kinase activity in a dose-dependent manner, as indicated by decreased phosphorylation of its substrate Rab10 at threonine 73 ^6^, as well as reduced LRRK2 S935 phosphorylation (Fig. 1H). Thus, genetically and chemically, the data show that LRRK2 inhibition enhances basal mitophagy in cells, and GSK3357679A displayed a superior performance compared to other available LRRK2 kinase inhibitors.

### Mutation of LRRK2 in vivo alters mitophagy in specific cell populations within the brain

Given the effects of LRRK2 kinase activity on mitophagy *in vitro*, we next sought to use our mouse lines to investigate this *in vivo*. PD is primarily a neurodegenerative disorder, so we first explored mitophagy in the brain. We focussed on four cell populations: two neuronal populations linked to movement - dopaminergic (DA) neurons of the substantia nigra pars compacta (SNpc) and Purkinje neurons of the cerebellum; as well as in two glial cell populations - cortical microglia and cortical astrocytes. In midbrain, we identified SNpc DA neurons using tyrosine hydroxylase (TH) staining and found no difference in the number of DA neurons per field across the *Lrrk2* genotypes (Fig. 2A and B). These cells are the mouse equivalent of the human population of DA neurons that degenerate in PD and we had previously found that they undergo substantial mitophagy ^17^. Basal mitophagy was enhanced in the LRRK2 KO neurons compared to the WT, and although not statistically significant, mitophagy appeared reduced in DA neurons of LRRK2 G2019S KI mice compared to WT (Fig. 2A and C). This was similar to our earlier observations in MEFs and showed that the presence of LRRK2 can impact mitophagy in this clinically relevant population of neurons within the midbrain. To determine if this effect is typical of neurons in general, we investigated mitophagy in another neuronal population involved in motor control, the Purkinje neurons. These cells were identified in cerebellar sections using immunostaining against the calcium sensor Calbindin-D28k. These cells are rich in mitochondria and as shown previously ^20^, they also undergo significant mitophagy (Fig. 2D). Contrary to what we observed in SNpc DA neurons, no statistical difference in mitophagy in Purkinje cells was found between any group (Fig. 2E).

**Figure 2.**
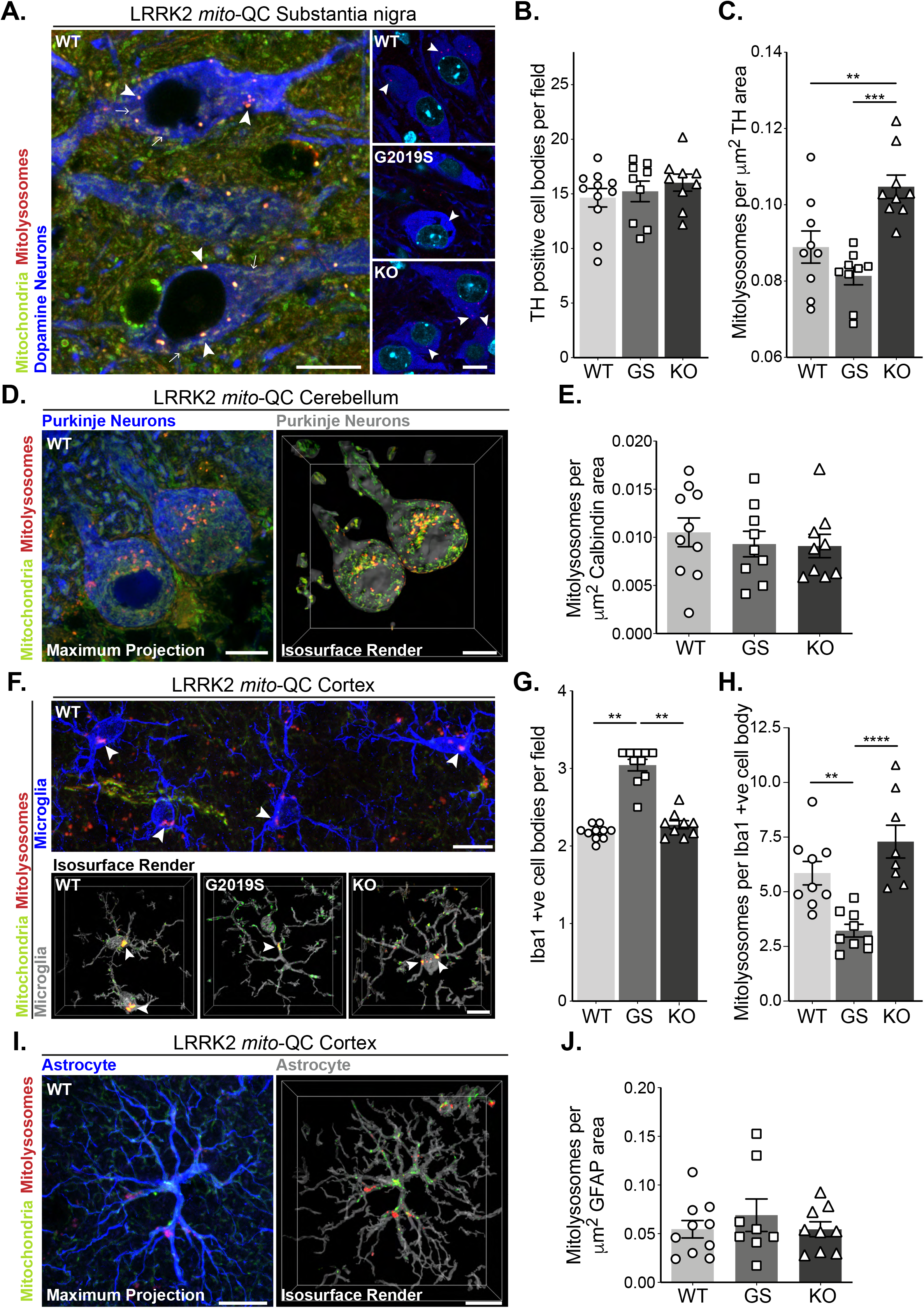
Mutation of LRRK2 *in vivo* alters brain mitophagy. (**A**) Representative image of tyrosine hydroxylase (TH) immunolabeled dopaminergic neurons within the substantia nigra pars compacta (SNpc) undergoing basal mitophagy in LRRK2 WT, LRRK2 G2019S, and LRRK2 KO *mito*-QC mice. Arrowheads show examples of mitolysosomes and arrows indicate mitochondria. (**B**) Quantitation of the number of TH positive cells per field of view using a 63x objective. (**C**) Quantitation of basal mitophagy per μm^2^ of TH staining, data points represent means from individual mice. (**D**) Representative maximal intensity projection and isosurface render of calbindin immunolabeled Purkinje neurons in *mito*-QC cerebellum sections, undergoing basal mitophagy. (**E**) Quantitation of basal mitophagy per μm^2^ of calbindin staining, data points represent means from individual mice. (**F**) Representative image and isosurface renders of Iba1 immunolabeled microglia undergoing basal mitophagy in cortical sections from LRRK2 WT, LRRK2 G2019S, and LRRK2 KO *mito*-QC mice. Arrowheads highlight mitolysosomes. (**G**) Quantitation of the number of Iba-1 positive cells in the brain cortex per field of view using a 63x objective. (**H**) Quantitation of basal mitophagy per Iba1 positive cell body, data points represent means from individual mice. (**I**) Representative maximal intensity projection and isosurface render of GFAP immunolabeled astrocytes undergoing basal mitophagy in *mito*-QC cortical sections. (**J**) Quantitation of basal mitophagy per μm^2^ of GFAP staining, data points represent means from individual mice. Scale bars, 10 μm. Overall data is represented as mean +/−SEM. Statistical significance is displayed as **p<0.01, and ***p<0.001.

As we observed neuron-specific alterations of mitophagy, we next examined the effect of *Lrrk2* genotype in two distinct populations glial cells within the cortex. Immune-related microglia were identified by Iba1 (ionized calcium-binding adapter molecule 1, Fig. 2F). Interestingly, we observed an enhanced presence of microglial cells in the cortex of G2019S animals when compared to WT or KO mice (Fig. 2G). We do not yet understand the nature of this increase and further work will be needed to determine if there are simply more microglia in the G2019S mice, or an increased movement of cells to this area of the brain. Regardless, when normalised for cell number, we found a significant decrease in basal mitophagy in G2019S microglia compared to WT, as well as an increase in mitophagy levels in KO cells (Fig. 2H). Thus, as with DA neurons, LRRK2 can impacts basal mitophagy in microglia. In contrast, cortical astrocytes, stained with glial fibrillary acidic protein (GFAP, Fig. 2I), did not show any observable difference in mitophagy across *Lrrk2* genotypes (Fig. 2J).

In contrast to the *Lrrk2* genotype effects on mitophagy in DA neurons and microglia, analysis of *auto*-QC mouse brains indicated no change in general macroautophagy in these cell types (Fig. S2). Thus, the LRRK2 G2019S mutation is not causing a major disruption in neuronal autophagy but does influence basal mitophagy levels.

### Mutation of LRRK2 in vivo also alters mitophagy in peripheral organs with high LRRK2 expression

We next assessed mitophagy levels in the lungs, a tissue in which the levels of LRRK2 are known to be elevated ^33,34^. Basal mitophagy across the whole lung was evident in all genotypes and in a similar fashion to MEFs, DA neurons and microglia, mitophagy was reduced in G2019S mice and enhanced (over 2-fold relative to WT) in KO mice (Fig. 3A and B), Consistent with this, mitochondrial content was decreased when comparing KO to G2019S (although no significant increase of this parameter was detected compared to WT, see Fig. S3A). As previously reported ^35,36^, we observed enlarged type II pneumocytes with large vesicular-like structures in all the animals of the LRRK2 KO group that is attributable to the accumulation of large lamellar bodies, which are secretory lysosomes responsible for surfactant release. We confirmed the nature of these structures as enlarged lamellar bodies by filipin staining lung sections for cholesterol, a component found in surfactant (Fig. S3B).

**Figure 3.**
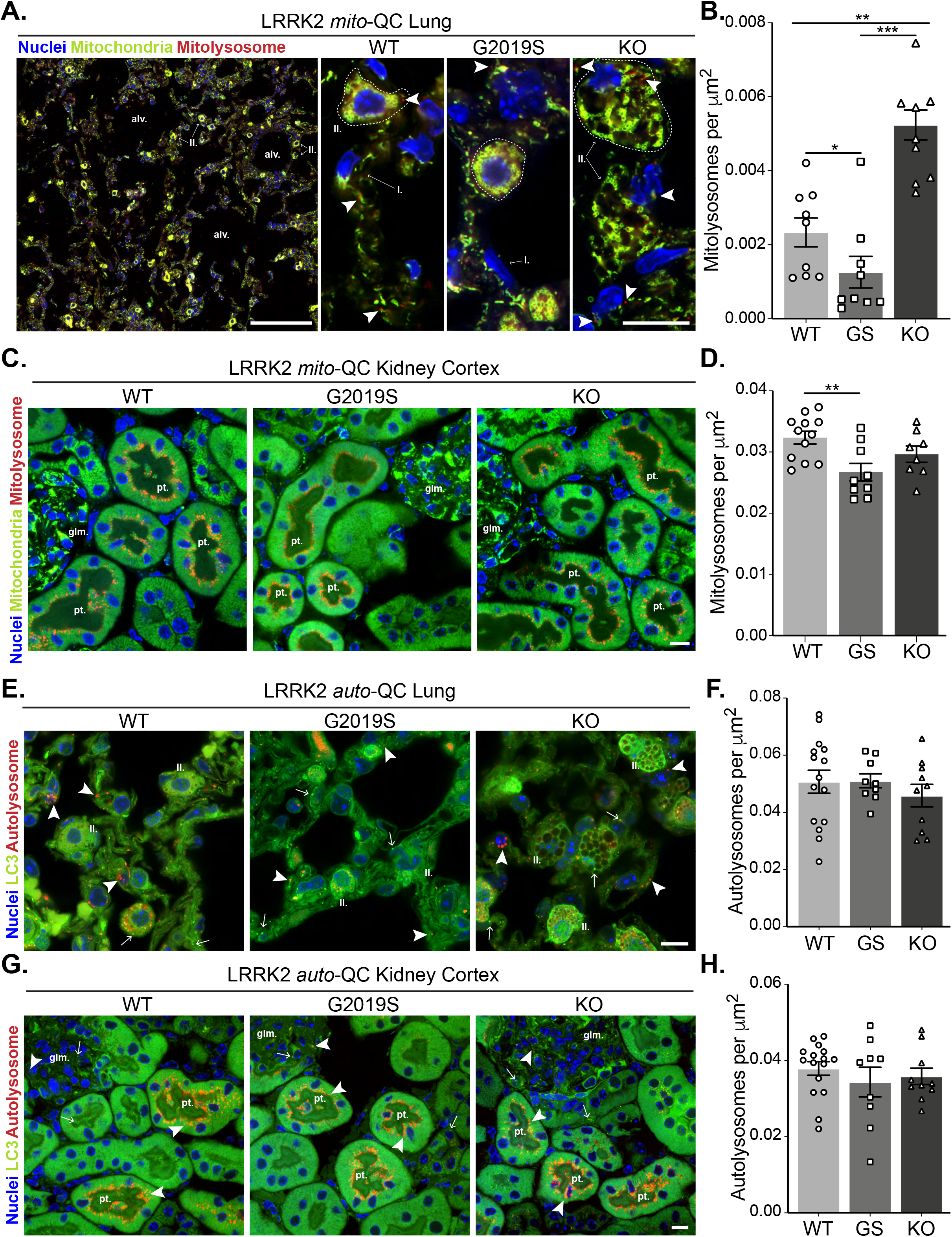
Effects of mutation of LRRK2 *in vivo* on basal mitophagy and macroautophagy in selected peripheral tissues. (**A**) Representative tile scan and images of *mito*-QC lungs from LRRK2 WT, LRRK2 G2019S, and LRRK2 KO mice. Arrows on the tile scan highlight Type II pneumocytes (II.), alv. show alveoli. Arrowheads in the higher magnification images indicate examples of mitolysosomes and circled cells correspond to Type II pneumocytes (II.) while I. indicates Type I pneumocytes. (**B**) Quantitation of lung basal mitophagy from data shown in D, data points represent means from individual mice. (**C**) Representative images of *mito*-QC kidney cortex from LRRK2 WT, LRRK2 G2019S, and LRRK2 KO mice. glm. indicates glomeruli and pt. indicates proximal tubule examples. (**D**) Quantitation of kidney basal mitophagy from data shown in F, data points represent means from individual mice. (**E**) Representative images of *auto*-QC lungs from LRRK2 WT, LRRK2 G2019S, and LRRK2 KO mice. Arrowheads highlight autolysosomes, arrows indicate autophagosomes, and II. indicates Type II pneumocytes. (**F**) Quantitation of lung autophagy from data shown in C, data points represent means from individual mice. (**G**) Representative images of *auto*-QC kidney cortex from LRRK2 WT, LRRK2 G2019S, and LRRK2 KO mice. Arrowheads and, arrows as in C, glm. indicates glomeruli, and pt. indicates proximal tubules. (**H**) Quantitation of kidney autophagy from data shown in E, data points represent means from individual mice. Scale bars, Tile scan in A: 100 μm, Other pictures:10 μm. Overall data is represented as mean +/−SEM. Statistical significance is displayed as **p<0.01, and ***p<0.001.

The tissue reported to have the highest LRRK2 expression is the kidney ^33,34^. We first investigated mitophagy in the kidney cortex, where we had previously shown the proximal tubules to be a major site of mammalian mitophagy ^20^. LRRK2-dependent mitophagy changes in the kidney were much lower in magnitude compared to the lung, yet there was a small decrease in G2019S-expressing tissue (Fig.3C and D). However, we do note that mitophagy is 10-fold higher in this region compared to lung, which may mask relatively small changes conferred by *Lrrk2* genotypes.

We next studied *in vivo* genotype effects on autophagy, using the same conditions and organs as for the *mito*-QC reporter. Consistent with brain and MEF data, no significant difference in the number of autolysosomes was observed in the lungs of *auto*-QC reporter mice (Fig. 3E and F). Again, enlarged type II pneumocytes were observed in LRRK2 KO animals (Fig. 3E). Likewise, in the kidney cortex we did not detect an effect of *LRRK2* genotype on autolysosomes (Fig. 3G and H). We also note that no major difference was seen in the number of autophagosomes across both lung and kidney (Fig. S3C and D). Taken together, these data suggest that the *Lrrk2* genotype does not majorly affect all autophagy pathways but predominantly impacts basal mitophagy, both *in vitro* and *in vivo*.

### GSK3357679A corrects the G2019S mitophagy defect in vivo

We next sought to determine if we could pharmacologically rescue the observed mitophagy defects *in vivo*. For this purpose, we utilised GSK3357679A - the pharmacodynamic characteristics of which have been shown to be suitable for extended oral dosing studies in rodents (Ding et al., in prep). We administered *mito*-QC WT, G2019S, and LRRK2 KO mice with GSK3357679A *via* oral gavage every 12 h for a total of four dosings. During this period, we observed no effect of GSK3357679A on body weight in any genotype (Fig. S4A). Tissues were then harvested 2 h post the final dosing. We focussed our analyses on tissues where our previous analyses of *Lrrk2* genotypic variants suggested a LRRK2-dependent role in mitophagy, namely the lung and brain as well as the kidney.

LRRK2 inhibition was confirmed in lung, brain and kidney tissue lysates of GSK3357679A dosed mice by immunoblotting of LRRK2 phopho-S935 and of Rab10 phospho-T73 (Fig. 4A). In the brain, GSK3357679A decreased the phosphorylation of LRRK2 on S935 in both WT and G2019S mice (Fig. 4A and B). However, we were not able to detect Rab10 phosphorylation in this tissue, suggesting Rab phosphorylation is low or tightly regulated (results not shown). Regardless, the loss of S935 phosphorylation on LRRK2 is consistent with kinase inhibition in the brain. LRRK2 kinase inhibition was also observed in the lungs and kidneys of the same animals, with GSK3357679A treated mice showing significant loss of both LRRK2 S935 phosphorylation and Rab 10 T73 phosphorylation (Fig. 4A and B).

**Figure 4.**
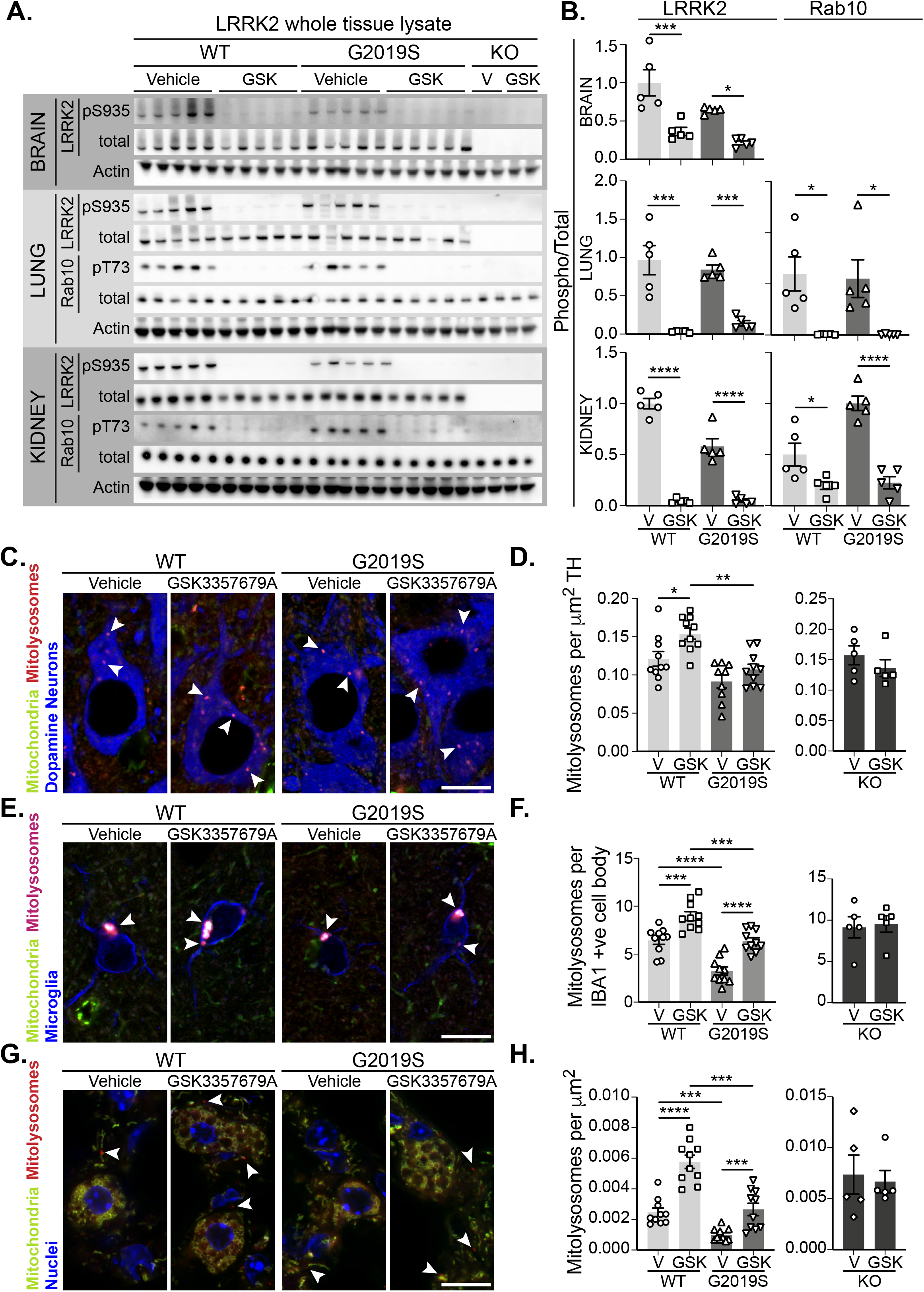
Pharmacological rescue of LRRK2-mediated mitophagy defects *in vivo.* (**A**) Immunoblots of the indicated proteins from tissue lysates of LRRK2 WT, LRRK2 G2019S, and LRRK2 KO *mito*-QC mice treated with vehicle or GSK3357679A. (**B**) Quantitation of phosphorylation data from A (**C**) Representative image of tyrosine hydroxylase (TH) immunolabeled dopaminergic neurons of the substantia nigra pars compacta in LRRK2 WT, LRRK2 G2019S, and LRRK2 KO *mito*-QC mice treated or not (vehicle) with GSK3357679A. Arrowheads indicate mitolysosome examples. (**D**) Quantitation of mitophagy from data shown in C, with the addition of LRRK2 KO. Each data point represents mean value from an individual mouse. (**E**) Representative images of Iba1 positive cortical microglia from LRRK2 WT, and LRRK2 G2019S mice treated or not (vehicle) with GSK3357679A. Arrowheads indicate mitolysosomes. (**F**) Quantitation of mitophagy from data shown in E, with the addition of LRRK2 KO. (**G**) Representative images of *mito*-QC lungs from LRRK2 WT, and LRRK2 G2019S mice treated or not (vehicle) with GSK3357679A. Arrowheads highlight mitolysosomes. (**H**) Quantitation of mitophagy from data shown in G, with the addition of LRRK2 KO. For (**B), (D), (F)**, and (**H)**, V = vehicle dosed animals, GSK = GSK3357679A dosed animals. Each data point represents mean value from an individual mouse. Scale bars, 10 μm. Overall data is represented as mean +/−SEM. Statistical significance is displayed as *p<0.05, **p<0.01, ***p<0.001, and ****p<0.0001.

In DA neurons of the SNpc, we found that treatment with GSK3357679A increased mitophagy in both WT and G2019S KI mice. Consistent with earlier results, the vehicle dosed G2019S group displayed significantly lower mitophagy compared to the vehicle- dosed WTs, (Fig. 4C and D, compare with Fig. 2A and C). Importantly, treatment with GSK3357679A restored G2019S mitophagy to base-line WT levels (Fig. 4C and D). No difference was observable in the KO groups in presence of GSK3357679A, showing mitophagy effects are through on-target LRRK2 inhibition.

As we had observed differences in cortical microglial mitophagy between genotypes, we next investigated the effect of GSK3357679A treatment on this cell population. GSK3357679A increased mitophagy in the cortical microglia in both WT and G2019S groups (Fig. 4E and F). Consistently, GSK3357679A restored G2019S mitophagy levels to a value similar to that observed in the control group (WT-V). Mitophagy was unaffected by GSK3357679A in the KO groups, confirming GSK3357679A specificity on LRRK2 kinase activity. As seen earlier (Fig. 2G), microglial cell numbers were increased in the cortex of vehicle-treated G2019S mice compared with vehicle-treated WT mice (Fig. S4B). Interestingly, GSK3357679A treatment recovered the increase in number of cortical microglia observed in LRRK2 G2019S KI mice (Fig S4B), suggesting that LRRK2 kinase activity is a key contributor to regulation of microglial numbers in this mouse line.

In the lungs we found that GSK3357679A increased mitophagy levels in both WT and G2019S KI animals (Fig. 4C and 4D). Importantly, in the G2019S group, GSK3357679A elevated mitophagy levels to a value similar to the WT Vehicle group, suggesting LRRK2-inhibition can rescue the G2019S-mediated defect in mitophagy in lung also. As observed for other LRRK2 inhibitors we observed enlarged lamellar bodies in Type-II pneumocytes in the lungs of mice treated with GSK3357679A, similar to that observed in LRRK2 KO mice (Fig. 3A) and to what has been previously reported in the presence of LRRK2 kinase inhibitors ^25,35,37^. GSK3357679A had no effect on mitophagy levels in the lungs of LRRK2 KO mice (Fig.4G). Consistent with the genetics, in the kidney we found that GSK3357679A had a minimal effect on mitophagy despite this organ exhibiting robust LRRK2 inhibition (Fig. S4B and C and Fig 4A). Though as previously mentioned, the very high levels of mitophagy in this tissue could be masking any subtle mitophagy increases.

Taken together these results show that a pathogenic mutation of LRRK2 impairs basal mitophagy in cells and tissues. Importantly, this phenotype can be rescued by the use of LRRK2 kinase inhibitors.

## Discussion

Our work reveals that LRRK2 kinase activity inversely correlates with basal mitophagy levels, both *in vitro* and *in vivo* in specific cells and tissues. Strikingly, the mitophagy defects seemed to be specific to certain cell types. Indeed, we observed different effects in two different neuronal subpopulations and in two microglial subpopulations. More work is needed to understand why mitophagy is more sensitive to LRRK2 kinase activity in these cells, but it may help explain why DA neurons degenerate in PD. Interestingly, we found a higher mitochondrial content in the soma of Purkinje neurons compared to DA neurons, indicative of a higher oxidative metabolism in the former. In addition, mitophagy levels were much lower in the Purkinje cells, implying an inverse correlation between oxidative metabolism and mitophagy, similar to previous observations in different muscle subtypes ^26^. Speculatively, the higher basal level of mitophagy in these DA neurons could be required to maintain oxidative metabolism in light of their lower mitochondrial numbers, thus rendering them susceptible to defects affecting mitophagy. Also of potential relevance, we observed a higher difference in mitophagy levels between genotypes in the cortical microglia than in the DA neurons. Brain resident microglia have been shown to have much higher LRRK2 levels and activity than neuronal populations, which could have implications for disease aetiology ^38^.

The work presented here shows for the first time that pathogenic LRRK2 mutations can alter basal mitophagy in clinically relevant cell populations *in vivo*. However, the extent to which impaired mitophagy drives an individual’s Parkinson’s disease remains to be determined. We and others have previously found that loss of PINK1 or Parkin activity does not significantly alter basal mitophagy rates *in vivo*, despite a well characterised role for the PINK1/Parkin pathway on depolarisation-induced mitophagy ^17,18,39^. This implies that the PINK1/Parkin pathway drives mitophagy under distinct types of stress, in contrast to the basally regulated LRRK2 pathway described here. Regardless, if loss of stress-induced PINK1/Parkin-dependent mitophagy can lead to PD, then it is reasonable to assume that loss of LRRK2-regulated basal mitophagy could also contribute. These data now imply that impaired mitophagy may be a common theme in PD pathology.

Recent reports in other models support our conclusion of a role for LRRK2 in regulating mitophagy. *In vitro* assays in patient derived fibroblasts bearing G2019S or R1441C LRRK2 variants are consistent with our *in vitro* cell assays and observations *in vivo* ^40–43^.

The mechanism by which LRRK2 kinase activity regulates basal mitophagy is currently unclear. LRRK2 has been shown to phosphorylate a subset of Rab GTPases ^6^ and, given the roles of Rabs in membrane trafficking, it is tempting to suggest that they may be key in regulating this mitophagy pathway ^8,44^. It has been recently shown that lysosomal overload stress induces translocation of Rab7L1 and LRRK2 to lysosomes ^45^. This leads to the activation of LRRK2 and the stabilisation of Rab8 and Rab10 through phosphorylation. Another recent study showed that LRRK2 mutations inhibit the mitochondrial localisation of Rab10 ^40^.

We used two highly similar reporter models in primary MEFs and in mice to study general autophagy and mitophagy. The use of both the *mito*-QC and the *auto*-QC reporters, in combination with selective LRRK2 kinase inhibitors, provided evidence that LRRK2 kinase activity affects mitophagy, rather than autophagy in general. The role of LRRK2 kinase activity on autophagy has been previously investigated in several studies with inconclusive or contradictory effects ^36,46–51^. However, with our reporter systems, we cannot entirely exclude that other selective autophagy pathways are affected. Additionally, total flux through the autophagy pathway is likely to be much higher than the relative flux attributable to mitophagy, and combined with the observed higher inter-individual variability with our autophagy reporter, this makes it potentially more difficult to pick up small changes. For these reasons, it would be unreasonable to entirely exclude the involvement of LRRK2 in general autophagy, although our results lack support for this.

Our results show that three structurally distinct selective LRRK2 kinase inhibitors are active on mitophagy in MEF cells, and that *in vivo* the tool compound GSK3357679A demonstrates similar cell-specific effects in DA neurons and microglia within the brain. Importantly, use of this inhibitor *in vitro* and *in vivo* supported our genetic data in suggesting that LRRK2 kinase activity inversely correlates with the level of basal mitophagy. The fact that we could rescue G2019S-impared mitophagy in PD-relevant cell types, within the brain, provides an exciting prospect that LRRK2 inhibitor-mediated correction of mitophagic defects in Parkinson’s patients could have therapeutic utility in the clinic. In addition, LRRK2 kinase activity inhibitors could also provide a way to increase mitophagy in general, which could be beneficial in idiopathic PD, or indeed, in other non-related conditions where increased clearance of mitochondria could be beneficial, such as mitochondrial diseases.

Here we demonstrate, through both genetic manipulation and pharmacology, that the most common mutation in PD impairs basal mitophagy in tissues and cells of clinical relevance. The fact that we can rescue this genetic defect in mitophagy using LRRK2 inhibitors, holds promise for future PD therapeutics.

## Acknowledgements

We would like to acknowledge Paul Appleton at the Dundee Imaging Facility, Dundee. The Zeiss LSM880 with Airyscan was supported by the ‘Wellcome Trust Multi-User Equipment Grant’ [208401/Z/17/Z]. We would also like to acknowledge Dr Jin-Feng Zhao and Dr Thomas McWilliams for their expert technical assistance. This work was funded by a grant from the Medical Research Council, UK (IGG; MC_UU_00018/2) and GlaxoSmithKline plc. Requests for provision of GSK3357679A should be directed to Alastair Reith.

## Author contribution

Conception and design were done by F.S., A.D.R. and I.G.G. Experiments were performed by F.S. and A.R.P. Data analysis was carried out by F.S. The autophagy counter plugin for FIJI was developed by G.B. and F.S. Drafting and revision of the manuscript was carried out by F.S., A.D.R. and I.G.G. All authors received and approved the final document.

---

**Figure S1.**
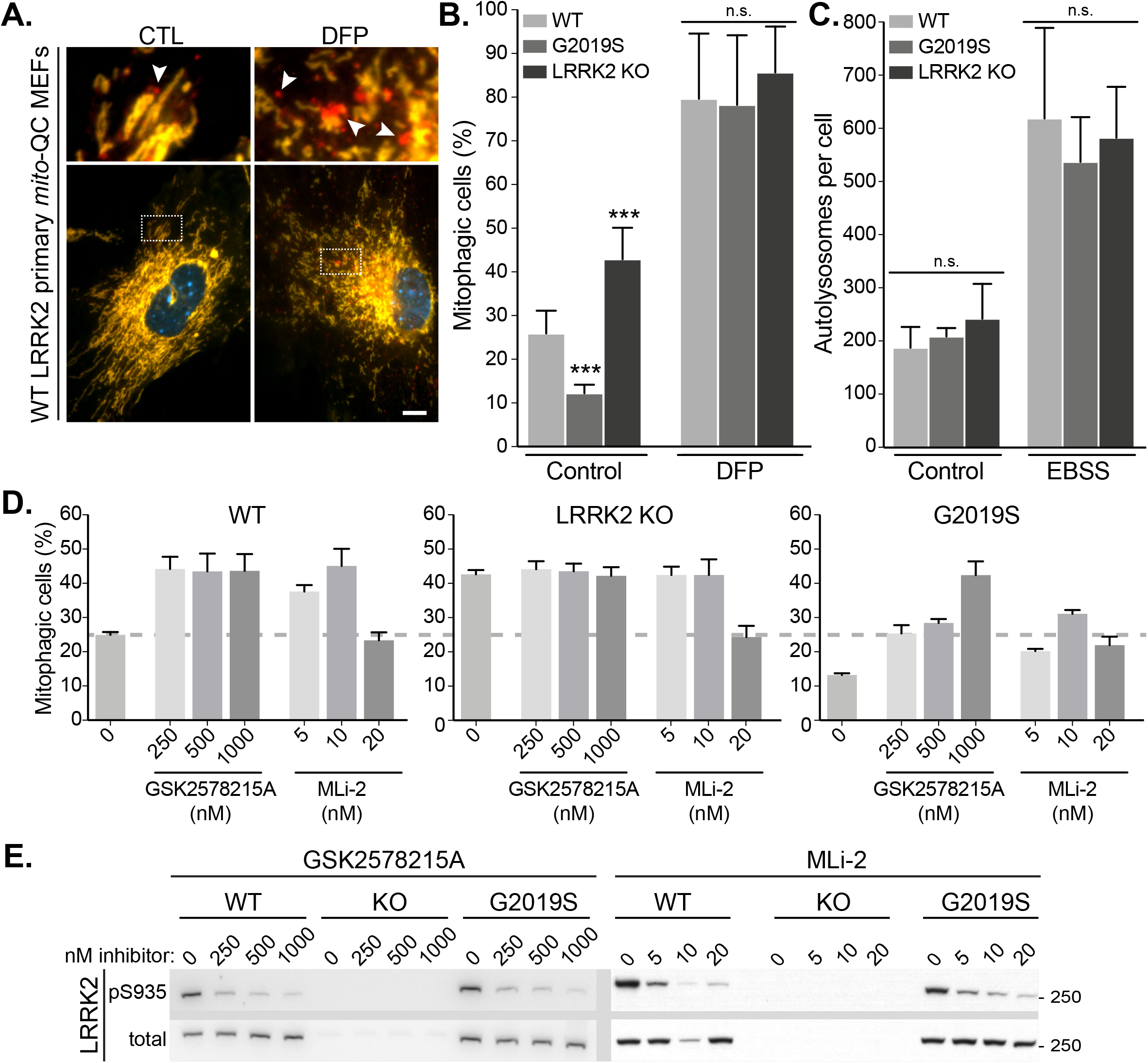
Stimulated mitophagy and autophagy are unchanged in MEFs. (**A**) Representative images of WT *mito*-QC primary MEF cultures treated with control (CTL) or 1mM deferiprone (DFP) for 24 h to stimulate mitophagy. Arrowheads indicate mitophagy (mCherry-only mitolysosomes). (**B**) Quantitation of basal (CTL) and stimulated (DFP) mitophagy in LRRK2 WT, LRRK2 G2019S, and LRRK2 KO *mito*-QC primary MEFs from 3-9 independent experiments. (**C**) Quantitation of *auto*-QC primary MEF cultures in basal (control) and amino acid starvation (EBSS) conditions from 4-6 independent experiments. (**D**) Quantitation of mitophagy in *mito*-QC MEFs, from 3-15 independent experiments treated with increasing concentrations of GSK2578215A and MLi-2. Data is represented as mean +/−SEM. Statistical significance is displayed as ***p<0.001. (**E**) Representative immunoblots MEFs treated as in D. Scale bars, 10 μm. Overall data is represented as mean +/−SEM. Statistical significance is displayed as ***p<0.001.

**Figure S2.**
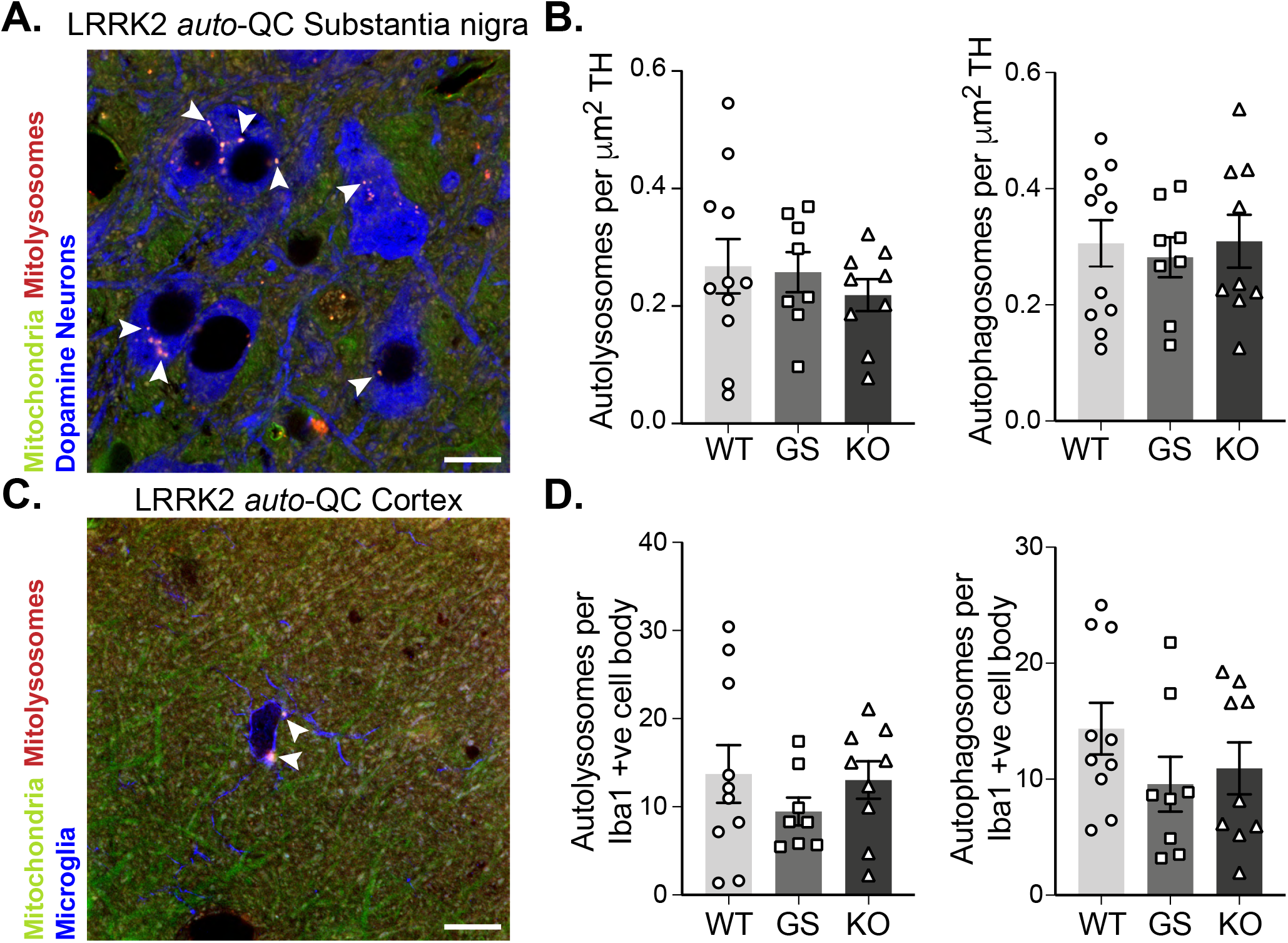
Macroautophagy in the brain is unaltered by *LRRK2* genotype. (**A**) Representative image of tyrosine hydroxylase (TH) immunolabeled dopaminergic neurons in substantia nigra pars compacta WT *auto*-QC sections. Arrowheads indicate autolysosome examples. (**B**) Quantitation of the number of autolysosomes (left) or autophagosomes (right) per TH area in LRRK2 WT, LRRK2 G2019S, and LRRK2 KO auto-QC mice. Data points are means from individual mice. (**C**) Representative image of Iba1 immunolabeled microglia in cortical WT *auto*-QC sections. Arrowheads indicate autolysosome examples. (**D**) Quantitation of the number of autolysosomes (left) or autophagosomes (right) per Iba1 positive cell in LRRK2 WT, LRRK2 G2019S, and LRRK2 KO *auto*-QC mice. Data points are means from individual mice. Scale bars, 10 μm. Overall data is represented as mean +/−SEM.

**Figure S3.**
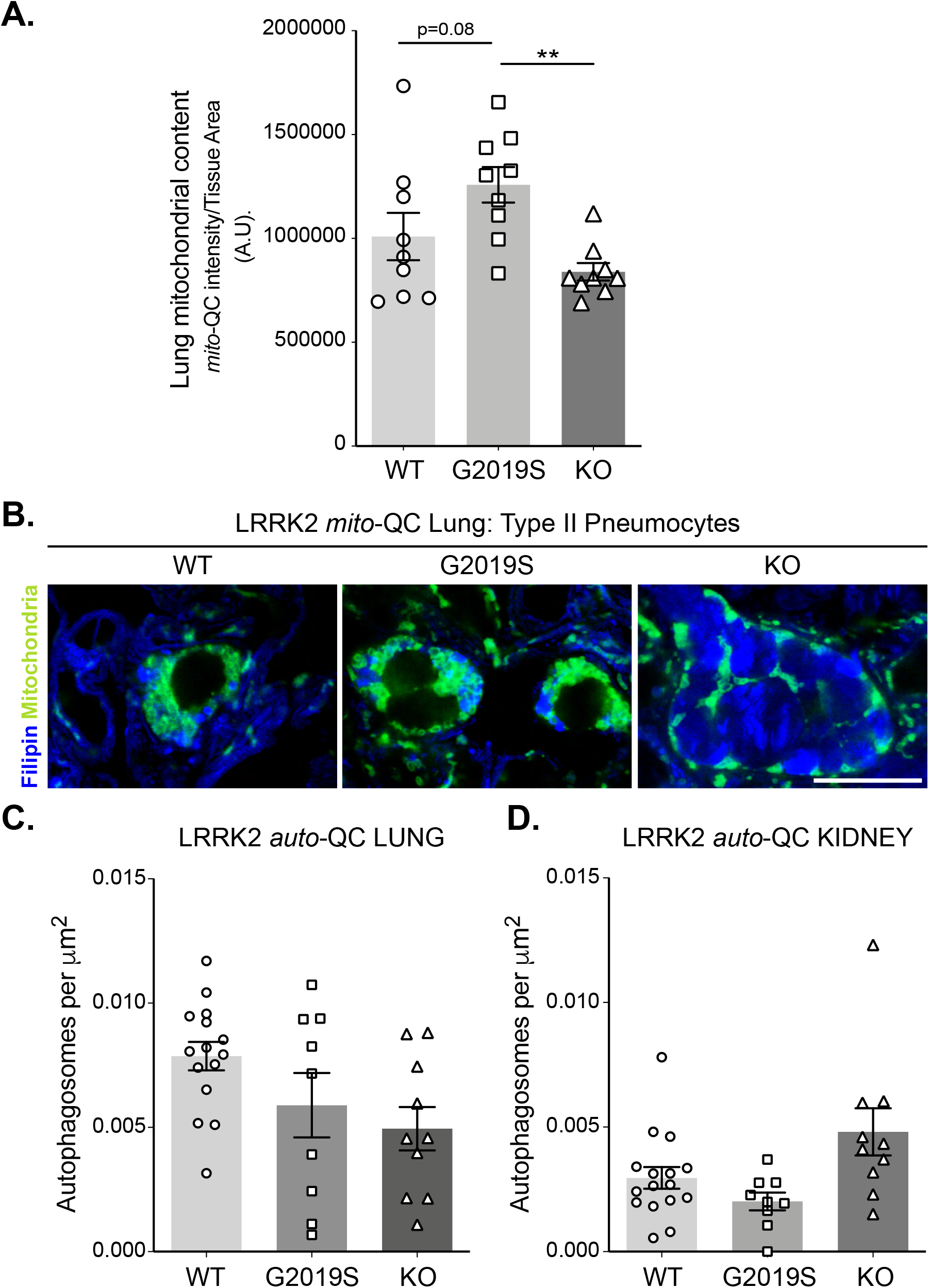
Macroautophagy in the lungs and kidney cortex is not changed by LRRK2 genotype. (**A**) Quantitation of mitochondrial content, based on total *mito*-QC reporter staining, from lung sections of indicated mice. Data points are means from individual animals. (**B**) Filipin staining of cholesterol in type II pneumocytes within *mito*-QC lungs. Note increased staining in KO. (**C**) Quantitation of basal number of autophagosomes in *auto*-QC lungs from LRRK2 WT, LRRK2 G2019S, and LRRK2 KO mice. (**D**) Quantitation of basal number of autophagosomes *auto*-QC kidney cortex from LRRK2 WT, LRRK2 G2019S, and LRRK2 KO mice. Scale bars, 10 μm. Data is represented as mean +/−SEM. Statistical significance is displayed as **p<0.01.

**Figure S4.**
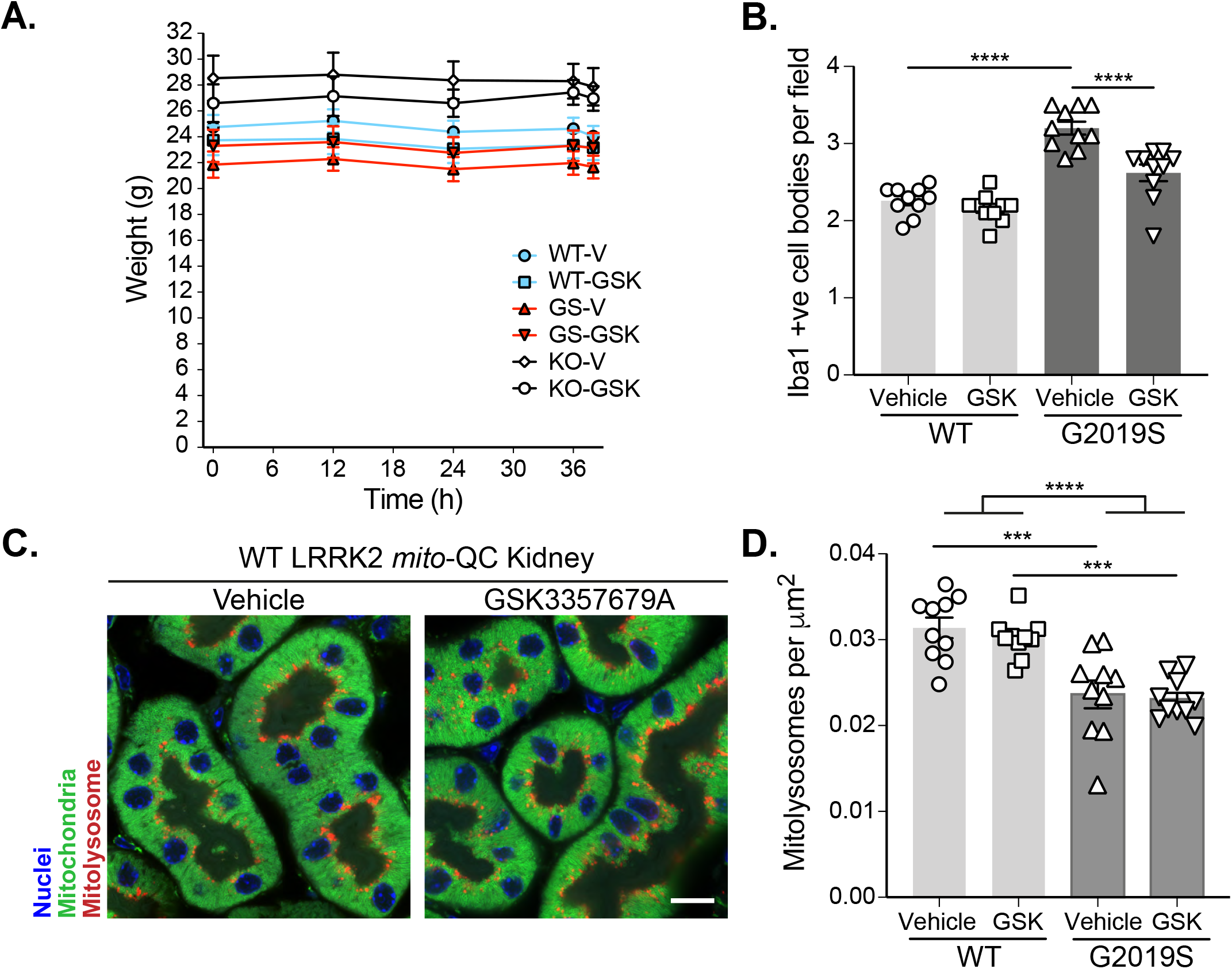
Pharmacological inhibition of LRRK2 kinase activity *in vitro* and *in vivo.* (**A**) Body weights of mice treated over 36 hours with vehicle (V) or GSK3357679A (X).(**B**) Quantitation of Iba-1 positive cells in the brain cortex per field of view (using a 63x objective) in LRRK2 WT, LRRK2 G2019S, and LRRK2 KO *mito*-QC mice treated with vehicle or GSK3357679A. (**C**) Representative images of *mito*-QC kidney cortex from LRRK2 WT mice treated with vehicle or GSK3357679A. (**D**) Quantitation of mitophagy from data shown in B. Data points represent the mean value from individual mice. Scale bars, 10 μm. Overall data is represented as mean +/−SEM. Statistical significance is displayed as ***p<0.001 and ****p<0.0001.

